# Frequent sugar feeding behavior by *Aedes aegypti* in Bamako, Mali makes them ideal candidates for control with attractive toxic sugar baits (ATSB)

**DOI:** 10.1101/574095

**Authors:** Fatoumata Sissoko, Amy Junnila, Mohamad M. Traore, Sekou F. Traore, Seydou Doumbia, Seydou Mamadou Dembele, Yosef Schlein, Petrányi Gergely, Rui-De Xue, Kristopher L. Arheart, Edita E. Revay, Vasiliy D. Kravchenko, John C. Beier, Gunter C. Müller

## Abstract

**Background:** Current tools and strategies are not sufficient to reliably address threats and outbreaks of arboviruses including Zika, dengue, chikungunya, and yellow fever. Hence there is a growing public health challenge to identify the best new control tools to use against the vector *Aedes aegypti*. In this study, we investigated *Ae. aegypti* sugar feeding strategies in Bamako, Mali, to determine if this species can be controlled effectively using attractive toxic sugar baits (ATSB).

**Methodology/Principal findings:** We determined the relative attraction of *Ae. aegypti* males and females to a variety of sugar sources including flowers, fruits, seedpods, and honeydew in the laboratory and using plant-baited traps in the field. Next, we observed the rhythm of blood feeding versus sugar feeding activity of *Ae. aegypti* in vegetation and in open areas. Finally, we studied the effectiveness of spraying vegetation with ATSB on *Ae. aegypti* in sugar rich (lush vegetation) and in sugar poor (sparse vegetation) urban environments.

Male and female laboratory sugar feeding rates within 24 h, on 8 of 16 plants offered were over 80%. The survival rates of mosquitoes on several plant sources were nearly as long as that of controls maintained on sucrose solution. In the field, females were highly attracted to 11 of 20 sugar sources, and 8 of these were attractive to males. Peak periods of host attraction for blood-feeding and sugar feeding in open areas were nearly identical and occurred shortly after sunrise and around sunset. In shaded areas, the first sugar-seeking peak occurred between 11:30 and 12:30 while the second was from 16:30 to 17:30. In a 50-day field trial, ATSB significantly reduced mean numbers of landing / biting female *Ae. aegypti* in the two types of vegetation. At sugar poor sites, the mean pre-treatment catch of 20.51 females on day 14 was reduced 70-fold to 0.29 on day 50. At sugar rich sites, the mean pre-treatment catch of 32.46 females on day 14 was reduced 10-fold to a mean of 3.20 females on day 50.

**Conclusions/Significance:** This is the first study to show how the vector *Ae. aegypti* depends on environmental resources of sugar for feeding and survival. The demonstration that *Ae. aegypti* populations rapidly collapsed after ATSB treatment, in both sugar rich and sugar poor environments, is strong evidence that *Ae. aegypti* is sugar-feeding frequently. Indeed, this study clearly demonstrates that *Ae. aegypti* mosquitoes depend on natural sugar resources, and a promising new method for vector control, ATSB, can be highly effective in the fight against Aedes-transmitted diseases.

**Author summary:** *Aedes aegypti* are notoriously difficult to control since their ubiquitous man-made and natural breeding sites, in various geographical regions, include almost any receptacle that can hold water. These diurnal mosquitoes are anthropophilic, a preference that promotes their role as vectors of many arboviruses including Zika, dengue, chikungunya, and yellow fever. With the exception of yellow fever, there are no vaccines against any of these arboviruses so that use of personal protective measures and mosquito vector control are the only means of prevention. Disease burdens in most endemic areas are not sufficiently reduced by various integrated vector management (IVM) strategies, hence there is a need for new control tools to complement the common strategies. Control by Attractive Toxic Sugar Baits (ATSB) appears to be an ideal candidate for this purpose.

The results of this study support this proposition. They demonstrate that *Ae. aegypti* in their urban environments in Mali are attracted to and frequently feed on staple diet that includes a variety of flowers, fruits and seed pods. Therefore, *Ae. aegypti* is a suitable candidate for control with ATSB. Moreover, the experiments with ATSB, in sparse vegetation or with competitor plant attractants in rich vegetation, demonstrated that ATSB treatment can cause a drastic reduction of *Ae. aegypti* populations.

## Introduction

*Aedes aegypti* are vectors for several significant arboviruses including Zika, dengue, chikungunya, and yellow fever [1-4]. Apart from yellow fever, there are no approved vaccines in widespread use against these viruses, therefore public health efforts are reliant on effective vector control methods [5,6]. Common strategies for the control of *Ae. aegypti* include residual spraying targeted to resting sites [7], space spraying indoors [8], larval control with conventional pesticides or lethal ovi-traps [9,10], and the use of personal protective measures.

However, the use of these methods is insufficient for achieving significant reductions in disease burdens thus new control tools are required [11,12]. Methods that are considered promising for this purpose by the WHO and require further field testing include: the Incompatible Insect Technique, the sterile insect technique, vector traps, and attractive toxic sugar baits [13, 14].

For many years, based on published work and experience in rearing laboratory mosquitoes, it was common knowledge that mosquitoes must feed on sugar to obtain the energy necessary for survival [15, 16]. However, the observation that *Ae. aegypti* could take blood and convert it to triglycerides and glycogen that provide energy [17] led to subsequent observations that female *Ae. aegypti* in the field were taking multiple blood meals within a single gonotrophic cycle [18-20]. Accordingly, *Ae. aegypti* may be less dependent on sugar for energy than other mosquito species and presumably rarely feed on sugar [18, 21, 22]. This is not necessarily a comprehensive rule since there are observations that *Ae. aegypti* are frequent sugar feeders, given the right environment. It was demonstrated that in Chiapas, Mexico, in an area with flowering plants, *Ae. aegypti* commonly fed on nectars; 8 to 21% of the mosquitoes tested positive for sugar using the anthrone test [23]. In the laboratory [24], it was observed that male and female *Ae. aegypti* displayed two peaks in sugar feeding; a small morning peak with 16 to 18% of mosquitoes sugar-feeding and a larger evening peak with 40 to 42% sugar-feeding mosquitoes. In another study in Duran, Ecuador, 56.8% of the *Ae. aegypti* females were marked as sugar positive following feeding on sucrose solution colored with food dye [25]. It has been concluded that the availability of sugar sources in the local environment affects mosquito longevity and thus it is a key regulator of mosquito population dynamics and therefore, of their vectorial capacity [15, 26]. The evidence above indicates that that the success of *Ae. aegypti*, at least in some regions, also depends on the presence of natural sugar sources. Attractive toxic sugar bait methods (ATSB) use the staple sugar feeding behavior of mosquitoes in nature for their control. The potential and efficiency of this method have been demonstrated in several experiments with various mosquito genera and species [27-34]. The observations on the sugar feeding behavior of *Ae. aegypti* indicate the potential of also using ATSB methods for the control of this species. The initial requirement for assessing the possibility of using ATSB against *Ae. aegypti* is the study of the sugar feeding behavior of the species on natural sugar sources in the experimental region. This was the basic purpose of this study which was investigated in the following manner: 1) in the laboratory, we observed whether *Ae. aegypti* groups were feeding on potentially edible sources of sugar, including flowers, fruits, seedpods, and honeydew, both available in the mosquito’s environment and some known to be highly attractive to other species; 2) We repeatedly offered single plant-sugar sources to mosquitoes as exclusive diets, one species per series of mosquitoes, and monitored the mortality rate in the mosquitoes; 3) We used plant baited traps in the field in and around the urban centers of Bamako, Mali and estimated plant attraction by comparing their catches of mosquitoes; 4) We determined the proportion of sugar fed mosquitoes and estimated sugar quantities in *Ae. aegypti* males and females from sugar rich and sugar poor urban neighborhoods; and 5) Finally, we tested the effect of ATSB treatment, estimated by the number of mosquitoes landing on volunteers, on *Ae. aegypti* population size in sugar rich and sugar poor areas in Bamako.

## Methods

### Identification of plants that are potential sugar sources for *Ae. aegypti*

Dominant plants at the study sites were defined with the assistance of the staff of the department of Traditional Medicine, School of Medicine and Dentistry, University of Sciences and Technology of Bamako, Mali.

### Mosquitoes for laboratory experiments

The *Ae. aegypti* colony used in the laboratory trials were established from the F1 generation of field caught mosquitoes to simulate the field breed as closely as possible. They were caught close to the campus of the campus of the University of Sciences, Techniques and Technology of Bamako, Mali, which is inside the city, and kept at their insectary under the following conditions: 27 ±3°C, relative humidity 70 ±10%, and photoperiod 12:12 hours light:dark. Adults for experiments were maintained on 10% sucrose solution and then starved for 24 h before the beginning of experiments. Fresh batches of 5 day old mosquitoes were used for each experiment.

### Testing for sugar

The sugar content in the gut of mosquitoes was determined by a modified cold anthrone test for fructose [35] and sugar amounts were visually scored by the intensity of the blue colour reaction under a dissecting microscope. Very light blue colour was defined as class I; darker blue colour in the reaction fluid was class II; class III was darker, and very intense blue was class IV (Fig 1).

**Fig 1.**
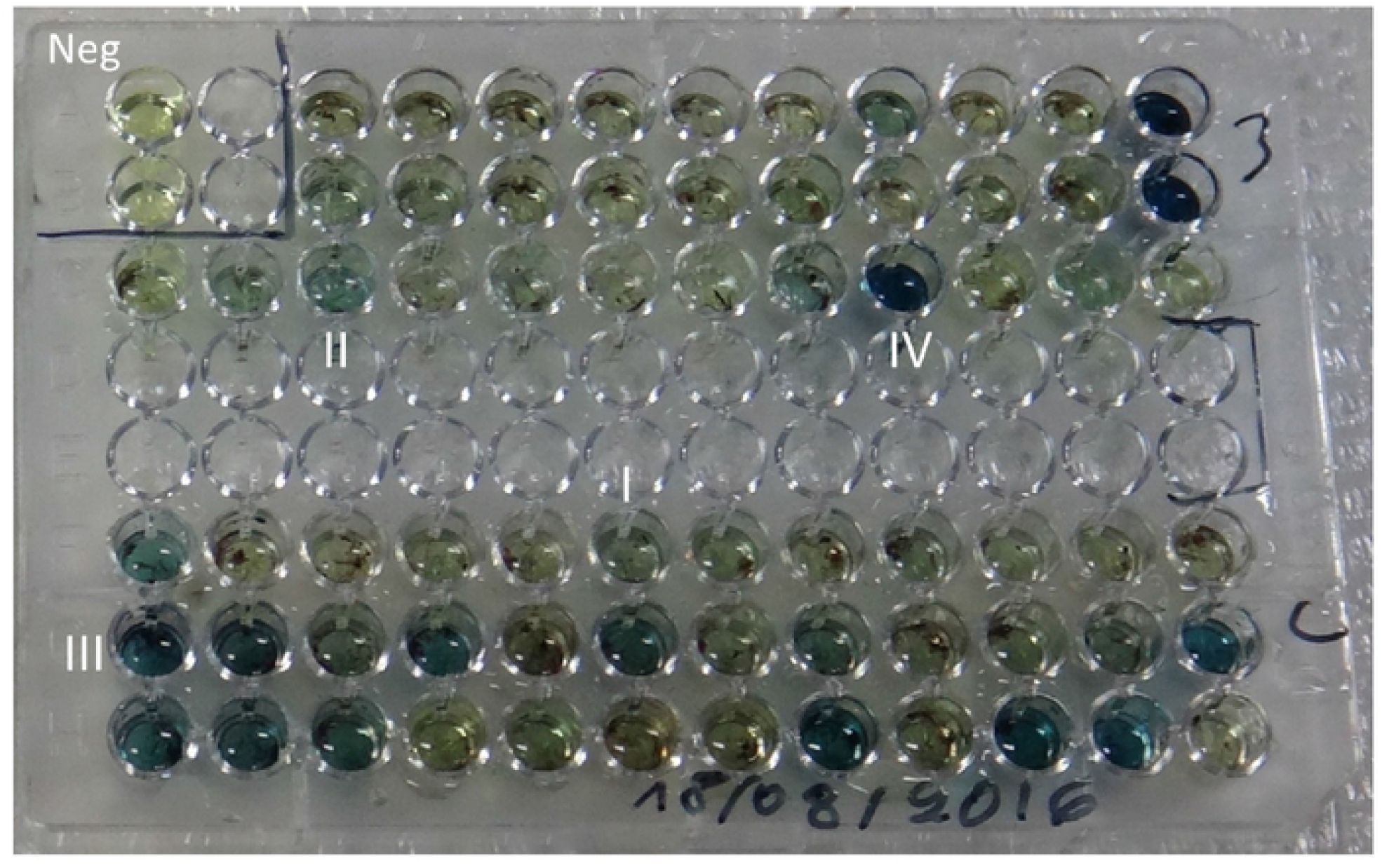
Microtitre plate with anthrone tested samples. Class I through IV colour intensities as well as negative control wells are shown.

### Laboratory experiments

#### Sugar feeding of *Ae. aegypti* exposed to potential sugar sources for 24 h

Assays were carried out with cohorts of 30 female and 30 male *Ae. aegypti* placed together for 24 h in 50 × 50 cm^2^ mosquito cages that contained one of the following: fruits of *Carica papaya* (papaya), *Mangifera indica* (mango), *Cucumis melo* (honeydew melon), leaves with extra-floral nectar of *Ricinus communis*, branches of *Lantana camara* soiled with honeydew from the aphid *Aphis gossypii*, and flowering *Prosopis juliflora*, *Acacia macrostachya*, *Acacia salicina*, *Lantana camara, Galphimia gracilis* and *Bougainvillia glabra*. Other types of offered sugar source were undamaged seedpods of *Piliostigma reticulatum*, infested *P. reticulatum* seedpods perforated by an unknown species of *Microlepidoptera* that were oozing sweet liquid, cut 20 cm segments of *Saccharum officinarum* (sugar cane) with the ends tightly sealed with parafilm, and broken / crushed tissue of similar segments of sugar cane. The general pattern was to offer either 4-5 branches of a target plant with stems in a beaker of distilled water, cut pieces of fruit (100 g), or complete seed pods (100 g). Additional tests were conducted on *Lantana camara* branches coated with ATSB solution (active ingredient: microencapsulated garlic oil) and ATSB offered in bait stations (active ingredient: dinotefuran). The bait stations were experimental prototypes with a thin protective membrane cover and were supplied from an ongoing Innovative Vector Control Consortium (IVCC) project carried out by Westham Ltd., Tel Aviv Israel.

A 10% sucrose solution and water soaked-cotton swabs served as the control diet and were made available in all cages. Each experiment was repeated 6 consecutive times. Mosquitoes were killed with CO_2_ and then either tested immediately for sugar by modified anthrone test or frozen at –70°C until anthrone tests could be performed.

#### Survival of *Ae. aegypti* females with continuous access to different sugar sources

Groups of 100 *Ae. aegypti* females in 50 × 50 cm cages, were allowed to feed on only 1 type of plant and their survival was monitored for 31 days. Plants were replaced daily, and water-soaked cotton swabs were generally available. Dead mosquitoes were removed daily. The 11 test diets were extra-floral nectar on foliage of non-flowering *Ricinus communis*, flowers of *Bougainvillia glabra, Prosopis juliflora*, or *Galphimia gracilis*, fruits of *Mangifera indica* and *Cucumis melo*, seed pods of *Piliostigma reticulatum*, either undamaged or perforated by pests and oozing juice, intact or broken / crushed stems of *Saccharum officinarum*, and honeydew (*Aphis gossypii*) soiled branches of non-flowering *Lantana camara*. Controls received 10% sucrose solution and water or water only. Each experiment was repeated 6 consecutive times.

### Field experiments

#### Field sites

All studies took place in Bamako, the capital and largest city of Mali, with ~ 2 million inhabitants. The city spreads out on both banks of the River Niger and annual flooding limits building on the banks which are a patchwork of wetlands, parkland, agricultural fields and forested areas. Some of the richer neighbourhoods and hotel districts include buildings with parklands or small vegetable fields. Bamako has tropical annual wet and dry periods. The hottest months are March, April, and May (hottest average temperature 32.4 ^0^C). The average temperature in the coldest month (December) is 25.1°C. Total annual rainfall averages 1098.5 mm; the rainy period is May through September, with peak rain occurring in August/September. The driest periods are late October through April.

‘Sugar rich’ and ‘sugar poor’ sites for field studies were chosen around Bamako and were identified primarily by land use which was determined by scouting areas by foot with trained botanists who could estimate flowering vegetation cover. Urban areas with the lush vegetation of irrigated parkland and gardens containing ≥50% flowering vegetation, were defined as ‘sugar rich’ and densely populated urban neighborhoods with sparse vegetation containing < 5% flowering vegetation, were defined as ‘sugar poor’.

#### Attraction of *Ae. aegypti* to plant baited glue net traps (GNT’s) in the field

Experiments were carried out for 10 consecutive days at the end of the dry season in 2016 along the shady margins of a forest gallery parallel to the River Niger.

Specifically designed glue net traps (GNTs) [30], were used to test attraction to plants. Briefly, cut bottom halves of 1.5 L plastic bottles were set in the ground, with the margins protruding to the surface, and filled with water. Dark green, rigid plastic netting, with mesh size 0.8 cm x 0.2 cm, was cut into 70 × 70 cm squares, rolled into cylinders, and each was put vertically above one of the cut bottles with the water, fastened to the ground with pegs and fitted with mesh covers. About 0.5 kg of test plant material was fixed in the center of a cylinder that was then covered and coated externally with glue (Tangle Foot, Tel Aviv, Israel). The different plant baits are listed in Table 2A and 2B. Controls were water-soaked sponges, empty traps, or naked *L. camara* branches, sprayed with either 10% sucrose solution or ATSB solution used in this study. Mosquitoes caught in the glue were removed, counted, and stored in 70% ethanol for identification. Each morning of the 10 day monitoring period, the baits were replaced, cylinders were repainted with glue, and trap locations were rotated to avoid location bias.

**Table 1.** Mean numbers ±SE over replicate cages in six experiments, in descending order, of sugar positive mosquitoes after 24 h exposure to a single plant-sugar source.

| Females |  |  |  |  | Males |  |  |  |  |
| --- | --- | --- | --- | --- | --- | --- | --- | --- | --- |
| Species | Mean | ±SE | % Positive | *p | Species | Mean | ±SE | % Positive | *p |
| *C. melo* F | 28.17 | 2.17 | 93.89% | 0.621 | *C. papaya* F | 29.17 | 2.21 | 97.22% | 0.830 |
| *A. macrostachya* FL | 28.00 | 0.91 | 93.00% | 0.601 | <sup>§</sup>ATSB on *L. camara* G | 28.50 | 2.18 | 95.00% | N/A |
| *P. juliflora* FL | 27.83 | 0.70 | 92.78% | 0.700 | *P. juliflora* FL | 28.00 | 2.16 | 93.33% | 0.871 |
| <sup>§</sup>ATSB on *L. camara* G | 26.67 | 1.33 | 88.89% | N/A | *A. macrostachya* FL | 27.83 | 2.15 | 92.78% | 0.828 |
| *A. salicina* FL | 26.50 | 2.10 | 88.33% | 0.956 | *G. gracilis* FL | 26.67 | 2.11 | 88.89% | 0.547 |
| *G. gracilis* FL | 25.50 | 2.06 | 85.00% | 0.693 | *A. salicina* FL | 26.50 | 2.10 | 88.33% | 0.511 |
| *C. papaya* F | 25.33 | 2.06 | 84.44% | 0.652 | *M. indica* F | 25.33 | 2.06 | 84.44% | 0.293 |
| *P. reticulatum* (oozing juice) F | 24.67 | 2.03 | 82.22% | 0.496 | *C. melo* F | 25.17 | 2.05 | 83.89% | 0.268 |
| *M. indica* F | 24.50 | 2.02 | 81.67% | 0.460 | <sup>§</sup>Sucrose 10% on cotton | 24.16 | 2.01 | 85.56% | 0.147 |
| <sup>§</sup>Sucrose 10% on cotton | 24.17 | 2.01 | 80.56% | 0.393 | *P. reticulatum* (oozing juice) F | 24.00 | 2.00 | 80.00% | 0.132 |
| <sup>§</sup>ATSB Bait Station | 18.83 | 1.77 | 62.78% | 0.006 | *L. camara* FL | 18.65 | 1.76 | 62.22% | 0.010 |
| *L. camara* FL | 18.81 | 1.75 | 61.11% | 0.003 | *L. camara* with *A. gossypii* G, H | 13.96 | 1.53 | 46.67% | <0.001 |
| *L. camara* with *A. gossypii* G, H | 14.50 | 1.56 | 48.33% | <0.001 | *R. communis* G, E | 13.31 | 1.49 | 45.00% | <0.001 |
| *S. officinarum* (broken) G | 13.17 | 1.48 | 43.89% | <0.001 | <sup>§</sup>ATSB Bait Station | 12.97 | 1.47 | 43.33% | <0.001 |
| *R. communis* G, E | 4.67 | 0.99 | 15.88% | <0.001 | *S. officinarum* (broken) G | 11.29 | 1.37 | 37.78% | <0.001 |
| *B. glabra* FL | 4.62 | 0.88 | 15.56% | <0.001 | *B. glabra* FL | 3.23 | 0.73 | 11.11% | <0.001 |
| *P. reticulatum* (intact) F | 0.67 | 0.33 | 2.22% | <0.001 | *P. reticulatum* (intact) F | 0.58 | 0.31 | 2.22% | <0.001 |
| *N. glauca* FL | 0.50 | 0.29 | 1.67% | <0.001 | *N. glauca* FL | 0.32 | 0.23 | 1.11% | <0.001 |
| *S. officinarum* (intact) G | 0.33 | 0.24 | 1.11% | <0.001 | *S. officinarum* (intact) G | 0.25 | 0.21 | 0.56% | <0.001 |

\* Compared to positive control ATSB on *L. camara* branches
F: Fruit, seedpod
<sup>§</sup>Positive control
FL: Blossom
N/A Not applicable
G: Branches
H: Honeydew on branches (*Aphis gossypii*)
E: Extra-floral nectar

**Table 2.**
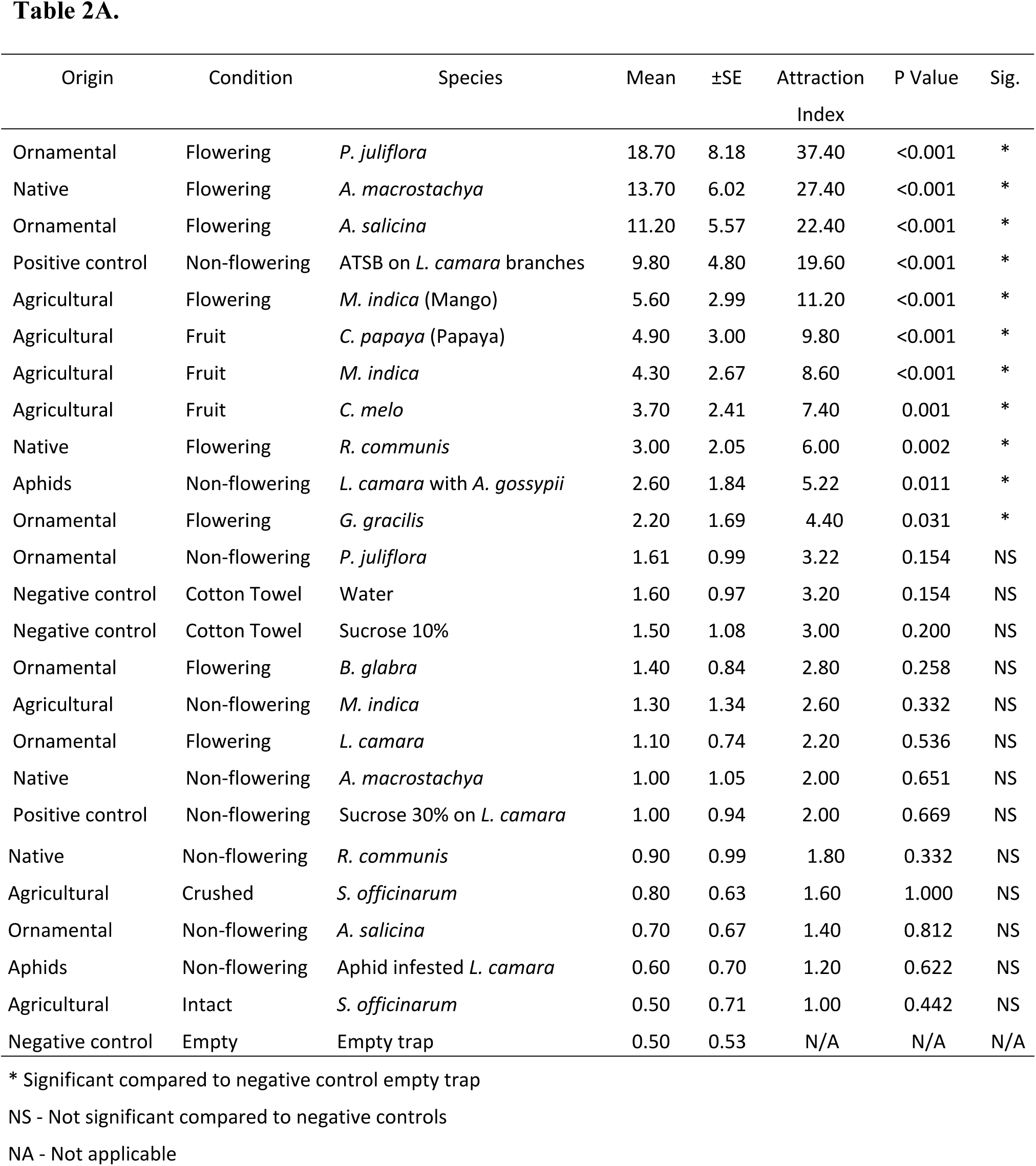
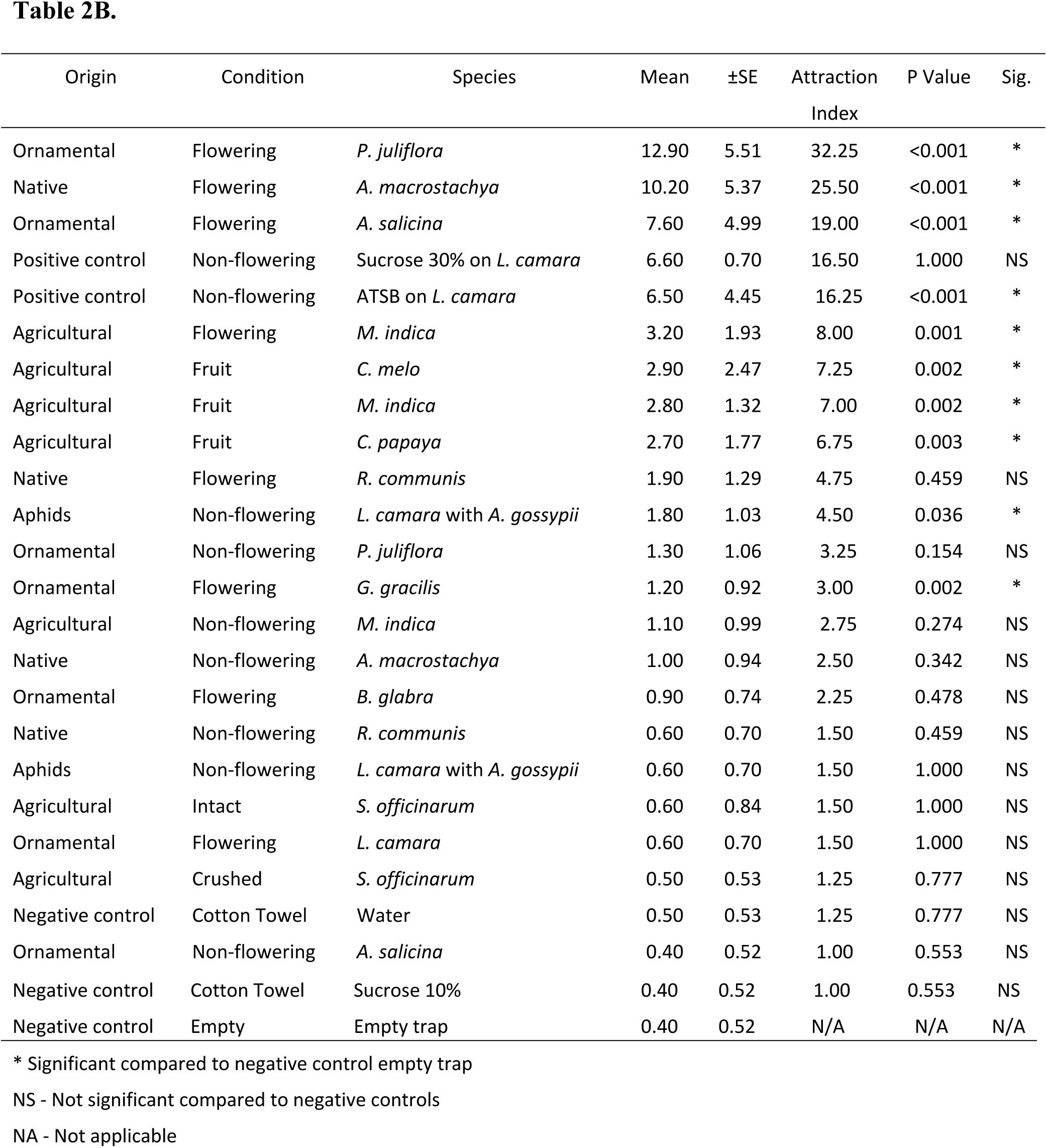
Mean number (± SE) of *Ae. aegypti* caught overnight by GNTs each with a different bait of fruit, seedpods, or flowers. **(A)** Females. **(B)** Males.

#### Timing of host-seeking and sugar-seeking activities in the field

The rhythm of the search for blood meals was shown by the number of landing/biting events on human volunteers in the field, in 30 min. intervals, for 18 hours (05:00 h to 23:00 h). This was done at 6 separate sites: 3 shady, 3 open and mosquitoes on volunteers from each group of sites were pooled and averaged. Mosquitoes were collected with aspirators near the shady margins of the forest gallery with thick undergrowth along the River Niger, and in open, sun-exposed grassland 30 m away from the trees. The United States Environmental Protection Agency guidelines and protocols for the use of human volunteers in landing catch experiments were carefully followed [36]. Three volunteers, 2 males and 1 female, all professional entomologists/medics participated in this study. As part of the consent process, the participants in all human trials were fully advised of the nature and objectives of the test and the potential health risks from exposure to mosquito bites. According to EPA regulations, they were required to avoid alcohol, caffeine, and fragrance products (e.g., perfume, cologne, hairspray, lotion, etc.) during the entire test period. For the tests, volunteers were wearing long trousers and long-sleeved shirts as protection against mosquito bites. One leg of the trousers was rolled up to expose the skin used as the test area. Volunteers were seated motionless in chairs with the exposed leg extended while observation, counting and recording of mosquitoes was made by assistants. The distance between the volunteers was 20 m and their locations were rotated through 9 stations every 30 min. to eliminate positional bias.

The rhythm of activity in the search for sugar meals was observed in both shaded and open areas by counting the catches of 9 GNTs in each type of area (18 total) baited with highly attractive flowering *P. juliflora* branches in 30 min. intervals, for 18 hours (05:00 h to 23:00 h). The baits were replaced by fresh branches at each time interval.

#### Comparing sugar feeding rates of *Ae. aegypti* in ‘sugar rich’ and ‘sugar poor’ habitats

The sugar-fed status of blood searching mosquitoes was observed. Catches on volunteers were carried out in the morning (11:00 h – 12:00 h) at ‘sugar poor’ and ‘sugar rich’ sites (Fig 2). Three volunteers, 2 males and 1 female, all professional entomologists/medics participated in this study and set-up was as described under “Timing of host seeking activity in the field”.

**Fig 2.**
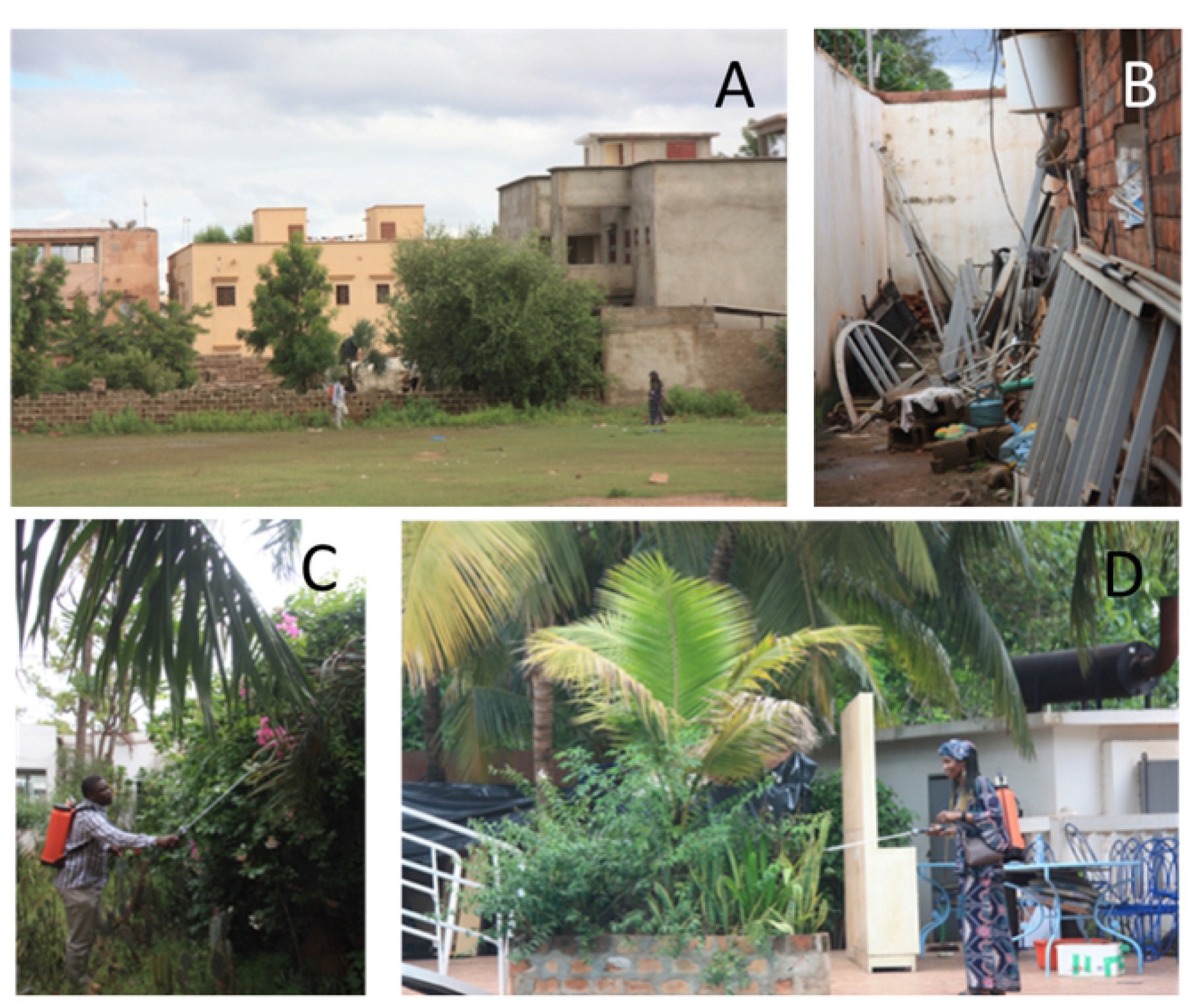
Pictures of sugar rich and sugar poor habitats in Bamako neighbourhoods. **A and B)** Densely populated urban areas with sparse vegetation (<5% flowering), **C and D**) Lush vegetation of irrigated gardens and parkland with areas of ≥50% flowering vegetation.

Sweep-net and aspirator (battery powered vacuum aspirators, John W. Hock, Gainesville, FL) catches for periods of 30 min. were also carried out in nearby vegetation. Captured mosquitoes were put on ice or in cages immediately, and random subsamples were tested for sugar [35].

Mosquito samples taken were either processed within 1 hour of collection or 6 hours after collection to assess the rate of sugar digestion from the guts of the mosquitoes. This process is expressed by the loss of positive anthrone reactions [35] with time. Random samples of 100 female and 100 male mosquitoes were tested from each site and for each processing time point. Sampling continued on consecutive days until these numbers were obtained, 4 days for the sugar rich area and 7 for the sugar poor area.

#### ATSB experiments

The ATSB method was field tested in 50 day experiments (Fig 2) in parklands and irrigated gardens (‘sugar rich’) and also in dry residential areas (‘sugar poor’) of Bamako from mid-June to the end of July 2016 (Fig 3).

**Fig 3.**
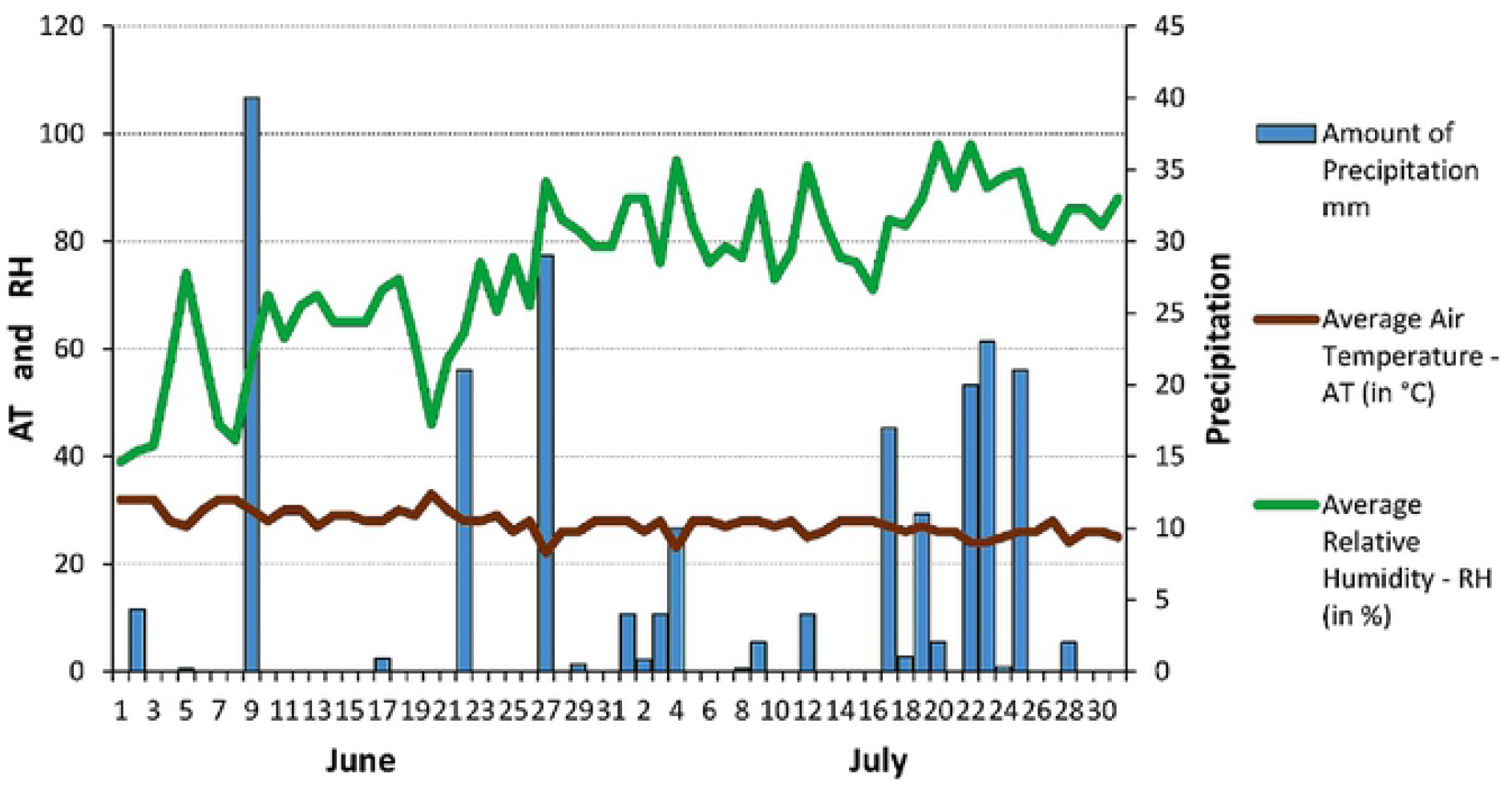
Local weather conditions in Bamako during June and July 2016. Rain occurred during the study, and two treatments were applied at the sites to avoid a “wash off’ effect.

A total of 12 sites, 6 ‘sugar rich’ and 6 ‘sugar poor’ areas of ~ 1 ha^2^ were selected. Three of each type were untreated controls and 3 in their vicinity received ATSB spray treatment. The effect of ATSB treatment was evaluated by recording biting / landing rates of female *Ae. aegypti* on human volunteers. The procedure was as described under “timing of host-seeking activity” with the same consent process [36]. The volunteers were placed in the center of each plot and each of them was moved among 3 monitoring stations per site that amounted to 9 repetitions per site.

ATSB was sprayed mainly on broad-leafed shrubs, bushes, and small trees up to 1.5 m high. Plants with flowers or fruit were not treated to minimize the impact on non-target organisms [37]. At the sugar poor sites, for the lack of suitable vegetation, artificial structures and buildings were sprayed. Treated areas were sprayed twice, once on day 15 and again on day 32 of the experiments.

#### ATSB formulations

For the ATSB spray trials, we used ATSB Mosquito Bait Concentrate, with the active ingredient of microencapsulated garlic oil (Universal Pest Solutions, Dallas TX, USA). The material was used according to label instructions, the concentrate was diluted 1:3, and was applied with a backpack sprayer on vegetation or on suitable artificial structures to cover 5% of the targeted area (four plots of 1 ha 500m^2^ were treated). Perimeter treatments were used at sites with continuous lines of vegetation (e.g. hedges, other shrubs) or extensive landscaping. ATSB was applied to vegetation in a continuous band measuring about 0.5 m wide, between 0.3 m to 1.5 m above the ground, and to the point of runoff at a rate of 500 to 600 ml of mixture per 30 m. Spot-treatments were applied to single shrubs, small patches of vegetation or artificial structures at least 30 cm above the ground in patches of approximately 1 m^2^ to the point of runoff. The spot applications were repeated, if possible, every 4 m.

### Statistical analysis

For the statistics reported in Table 1, we used a generalized linear model for a Poisson distributed outcome, number of female and male mosquitos. Separate models were employed for females and males. The independent variable was plant. We present model means and standard deviations along with raw and Dunnett adjusted p-values for comparing means with the control, ATSB on *L. camara* branches.

Figures 4 A and B present survival plots for a series of female *Ae. aegypti* that were each exposed for 31 days to two types of plants. We used Kaplan-Meyer statistics to compute the data for these figures.

**Fig 4.**
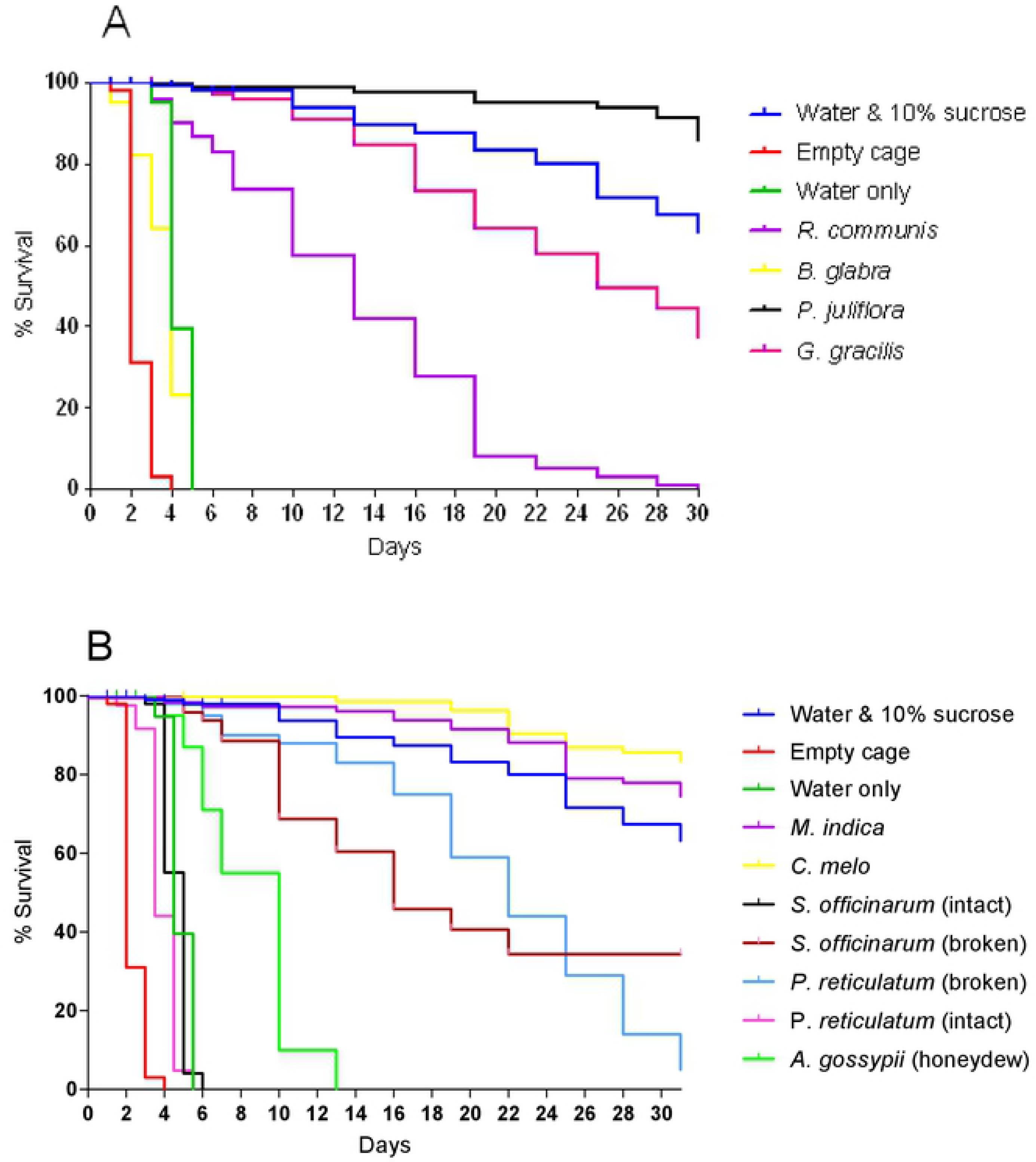
Survival in a series of female *Ae. aegypti* that were each exposed for 31 days to one type of plant. **(A)** Survival after exposure to branches with plant blossoms. *Ricinus communis* baits were branches with extra-floral nectaries. **(B)** Sugar source types other than blossoms.

Tables 2 A and B report the mean number of *Ae. aegypti* caught overnight by GNTs each with a different bait of fruit, seedpods, or flowers. We used a generalized linear model for a Poisson distributed outcome, number of female and male mosquitos. Overdispersion was evident; therefore, we changed to a negative binomial model that employs a scale parameter to adjust the model for overdispersion. Separate models were analyzed for females and males. The independent variable was plant. We present model means and standard deviations along with raw and Dunnett adjusted p-values for comparing means with the control, an empty trap. The data for Figures 4 A and B come from the data used in the analysis of Tables 2 A and B.

The percentage of *Ae. aegypti* with different amounts of sugar in the gut is reported in Table 3. The mosquitos were caught at ‘sugar rich’ and ‘sugar poor’ sites on human volunteers or with sweep-nets in the vegetation. We totaled the number of sugar positive mosquitos in each category as determined by anthrone testing for each time, sugar status, and catch type and then divided by the total number of sugar positive mosquitos. The data for Figures 5 A and B come from the data analyzed for Table 3.

**Table 3.** The percentage of *Ae. aegypti* with different amounts of sugar in the gut that were caught at ‘sugar rich’ and ‘sugar poor’ sites on human volunteers or with sweep-nets in the vegetation. Sugar quantities were classed by the intensity of the anthrone reaction.

| Site | Time<br>* | Catch<br>type | Females |  |  |  | Males |  |  |  |
| --- | --- | --- | --- | --- | --- | --- | --- | --- | --- | --- |
|  |  |  | I | II | III | IV | I | II | III | IV |
| Sugar rich | 1 hr | Human landing catch | 60.32 | 32.54 | 6.35 | 0.79 | 23.00 | 40.00 | 29.00 | 8.00 |
| Sugar poor | 1 hr | Human landing catch | 68.37 | 24.49 | 7.14 | 0.00 | 28.57 | 48.57 | 18.57 | 4.29 |
| Sugar rich | 1 hr | Resting site catch | 13.18 | 21.47 | 40.49 | 24.86 | 11.60 | 20.63 | 42.54 | 25.23 |
| Sugar poor | 1 hr | Resting site catch | 46.31 | 28.19 | 19.80 | 5.71 | 53.94 | 37.54 | 7.26 | 1.26 |
| Sugar rich | 6 hr | Human landing catch | 90.39 | 9.62 | 0.00 | 0.00 | 81.81 | 15.91 | 2.27 | 0.00 |
| Sugar poor | 6 hr | Human landing catch | 92.31 | 7.69 | 0.00 | 0.00 | 56.94 | 30.56 | 12.50 | 0.00 |
| Sugar rich | 6 hr | Resting site catch | 47.15 | 36.94 | 12.61 | 3.30 | 56.69 | 29.94 | 11.78 | 1.59 |
| Sugar poor | 6 hr | Resting site catch | 80.00 | 16.92 | 3.08 | 0.00 | 75.00 | 23.53 | 1.47 | 0.00 |

\*Time between collection and testing

**Fig 5.**
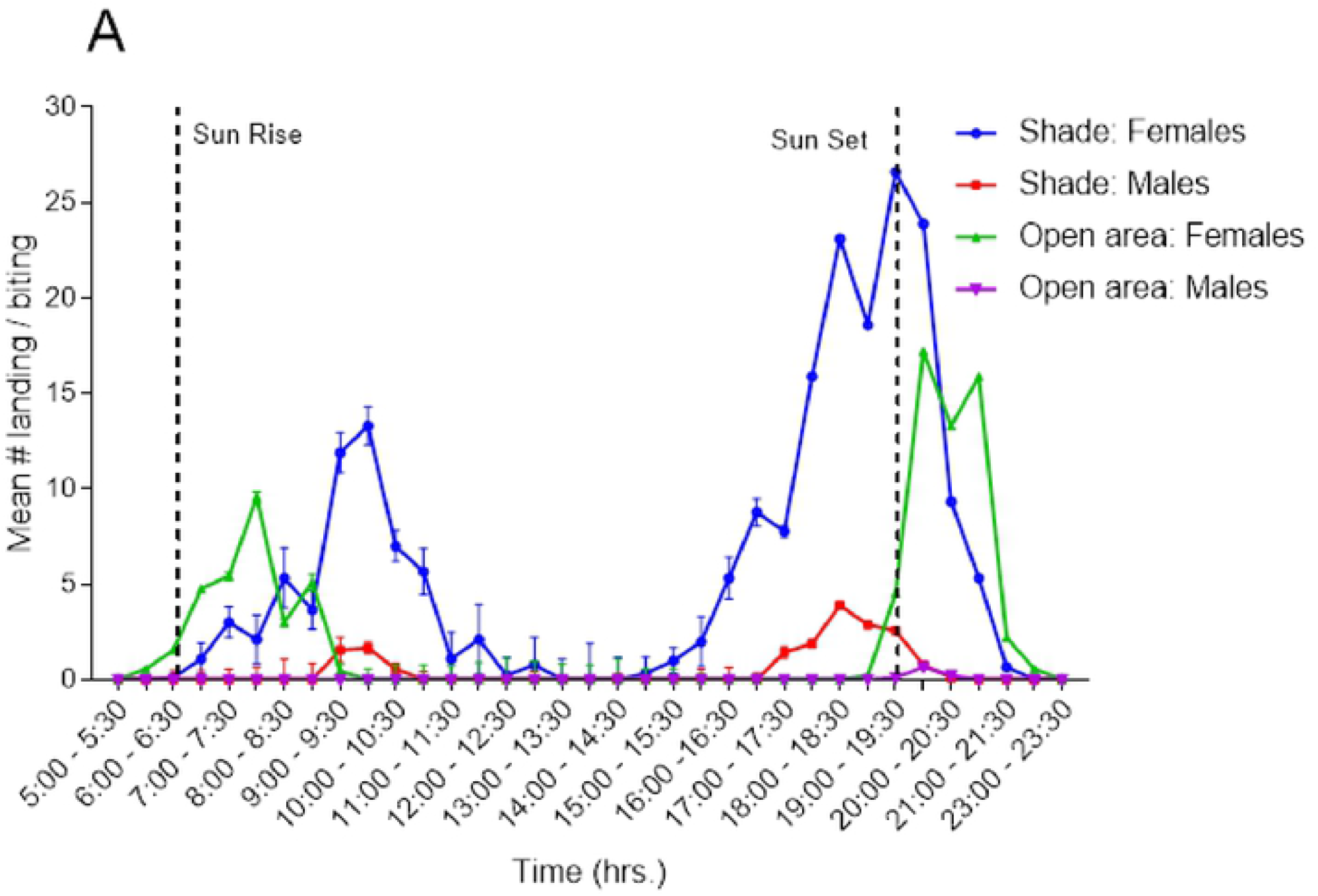
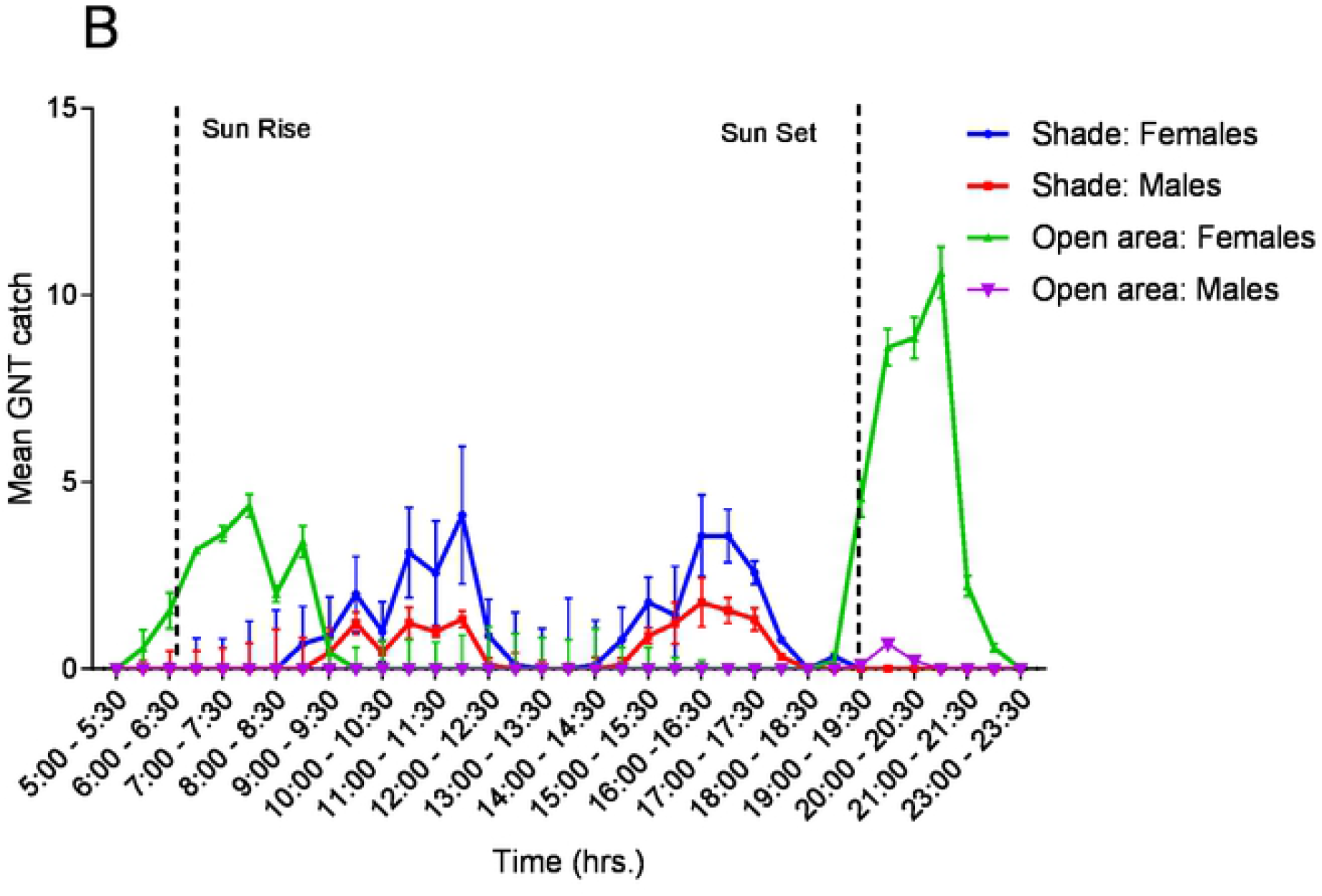
Periodicity of host-seeking and sugar-seeking behavior of *Ae. aegypti* over 18 hours. **(A)** evaluated by average catches of mosquitoes landing on a human volunteer, in 30 min intervals (±SE) and **(B)** evaluated by catches of GNTs baited with highly attractive flowering branches of *P. juliflora*. Numbers shown are the average catch per volunteer, per time period and average catch per trap, per time period (±SE).

Table 4 presents the mean number of *Ae. aegypti* following ATSB treatment at sugar poor and sugar rich sites. The data is the number of female mosquitos landing on human volunteers compared to their frequency at untreated control sites. We applied a generalized linear mixed model analysis to the mean catch for control and treatment period in each sugar rich and sugar poor control and experimental groups. Each mean came from nine catches: three volunteers each providing landing catches in three different monitoring stations. The Shapiro-Wilk test determined that the mean counts were normally distributed. The model included fixed effects for group (sugar rich and sugar poor), treatment period (control [days 1-14], ATSB treatment 1 [days 16-31], and ATSB treatment 2 [days 35-50]), condition (control and experimental), plus all two-way interactions and the three-way interaction. Appropriate error terms for the fixed effects are random effects. The error term for the group effect was site nested within group. The treatment and condition effects were repeated measures within the groups. Therefore, the error term for treatment and group x treatment was group x treatment x site nested within group. Likewise, the error term for condition and condition x group was condition x group x site nested within group. The error term for treatment x condition and group x treatment x condition was treatment x condition x site nested within group. Means and standard errors are reported for the group x treatment x condition interaction. We report p-values and Bonferroni adjusted p-values for comparisons between control and experimental means at each treatment level for sugar poor and sugar rich groups as well as comparisons between sugar rich and sugar poor means at each treatment level for control and experimental groups. Finally, we report p-values and Dunnett adjusted p-values for comparisons between control and the two ATSB levels for sugar rich and poor experimental and control groups. The data for this analysis was used to produce Figures 7 A and B.

**Table 4.** Reduction of the *Ae. aegypti* population following ATSB treatment at ‘sugar poor’ and ‘sugar rich’ sites as indicated by the decrease in landings of females on human volunteers compared to their frequency at untreated control sites.

| Day | Sugar Poor |  |  |  |  | Sugar Rich |  |  |  |  | SP vs. SR |  |
| --- | --- | --- | --- | --- | --- | --- | --- | --- | --- | --- | --- | --- |
|  | Control |  | Experimental |  |  | Control |  | Experimental |  |  | Cont | Exp |
|  | Mean | SE | Mean | SE | P | Mean | SE | Mean | SE | P | P | P |
| 1 | 5.46 | 0.63 | 9.10 | 0.94 | 0.001 | 12.27 | 1.20 | 19.41 | 1.78 | 0.001 | <0.001 | <0.001 |
| 3 | 8.51 | 0.89 | 8.98 | 0.93 | 0.716 | 13.69 | 1.32 | 17.53 | 1.63 | 0.065 | 0.001 | <0.001 |
| 6 | 6.66 | 0.73 | 10.17 | 1.03 | 0.005 | 19.68 | 1.80 | 20.39 | 1.86 | 0.785 | <0.001 | <0.001 |
| 9 | 7.99 | 0.84 | 12.96 | 1.26 | 0.001 | 16.00 | 1.50 | 22.94 | 2.07 | 0.006 | <0.001 | <0.001 |
| 12 | 14.05 | 1.35 | 16.67 | 1.56 | 0.203 | 25.70 | 2.29 | 23.62 | 2.12 | 0.505 | <0.001 | 0.007 |
| 14 | 16.46 | 1.55 | 20.51 | 1.88 | 0.096 | 22.67 | 2.05 | 32.46 | 2.84 | 0.004 | 0.014 | 0.000 |
| 16 | 19.19 | 1.77 | 8.58 | 0.90 | <0.001 | 24.59 | 2.21 | 22.77 | 2.06 | 0.548 | 0.054 | <0.001 |
| 19 | 21.04 | 1.93 | 4.81 | 0.58 | <0.001 | 28.21 | 2.50 | 17.46 | 1.62 | 0.000 | 0.022 | <0.001 |
| 22 | 17.06 | 1.59 | 2.49 | 0.36 | <0.001 | 25.17 | 2.25 | 10.31 | 1.04 | <0.001 | 0.003 | <0.001 |
| 25 | 23.28 | 2.10 | 1.56 | 0.27 | <0.001 | 30.99 | 2.73 | 7.60 | 0.81 | <0.001 | 0.024 | <0.001 |
| 28 | 18.76 | 1.73 | 0.70 | 0.17 | <0.001 | 38.10 | 3.31 | 3.95 | 0.50 | <0.001 | <0.001 | <0.001 |
| 31 | 23.57 | 2.12 | 0.47 | 0.14 | <0.001 | 44.29 | 3.82 | 6.62 | 0.73 | <0.001 | <0.001 | <0.001 |
| 35 | 26.04 | 2.32 | 0.22 | 0.09 | <0.001 | 40.65 | 3.53 | 2.44 | 0.36 | <0.001 | 0.000 | <0.001 |
| 38 | 24.34 | 2.19 | 0.41 | 0.13 | <0.001 | 48.95 | 4.18 | 1.14 | 0.22 | <0.001 | <0.001 | 0.005 |
| 41 | 28.89 | 2.55 | 0.26 | 0.10 | <0.001 | 55.37 | 4.69 | 0.92 | 0.20 | <0.001 | <0.001 | 0.004 |
| 44 | 24.23 | 2.18 | 0.63 | 0.16 | <0.001 | 62.17 | 5.24 | 1.00 | 0.21 | <0.001 | <0.001 | 0.156 |
| 47 | 30.14 | 2.66 | 0.47 | 0.14 | <0.001 | 56.52 | 4.78 | 0.59 | 0.16 | <0.001 | <0.001 | 0.561 |
| 50 | 32.93 | 2.89 | 0.29 | 0.10 | <0.001 | 65.89 | 5.55 | 3.20 | 0.43 | <0.001 | <0.001 | <0.001 |

Plants in field experiments were also ranked by being assigned an attraction index (AI), which was calculated by the following equation: average catch with the plant bait (PB) ÷ by the average catch with the empty trap control (ET); PB/ET = AI.

## Results

### Sugar feeding of *Ae. aegypti* exposed to potential sugar sources for 24 h

Sugar feeding rates for females on 8 out of 16 of the sugar sources were high, ranging from 81.67% on *Mangifera indica* (mango), to 93.89% on *Cucumis melo* (honeydew melon). Males fed avidly on 8 sugar sources. Exposure to fluids oozing from decomposing seedpods of *P. reticulatum* resulted in 80% sugar positive mosquitoes and the highest result was the 97.22% of positive specimens in the series that received *C. papaya* (papaya). Also, 88.89% of the females and 95.00% of the males in the ATSB on *L. camara* group were sugar positive after the exposure (Table 1).

### Survival of *Ae. aegypti* females with continuous access to different sugar sources

Of the different female groups fed exclusively for 31 days on one diet, the negative control group of 100 starved and thirsty females died within 4 days. Mosquitoes survived for up to 6 days on water alone and the provision of 10% sucrose solution allowed 68% survival of up to 31 days. Among the plant diet series, the survival proportion by day 31 was the highest (85.89%) in the group fed on *P. juliflora* whereas the lowest survival rate of 5.00% was in the group that received intact seedpods of *P. reticulatum* (Fig 4A and 4B).

### Attraction of *Ae. aegypti* to plant baited glue net traps (GNTs) in the field

Females were attracted to 11 of the 23 baits. The highest mean catches of 18.70, 13.70, 11.20, and 9.80 specimens were from traps baited with *P. juliflora, A. macrostachya, A. salicina, M. indica*, and ATSB on branches, respectively. Males were attracted to 8 baits, the 4 most attractive being *P. juliflora* (12.90), *A. macrostachya* (10.20), *A. salicina* (7.60), and the positive control of ATSB coated branches (6.50) (Tables 2A and 2B).

### Timing of host-seeking and sugar-seeking activities in the field

In shady areas, the three volunteers caught an average 54.5 females and 4.6 males over 18 hours of monitoring. The same volunteers in an open area caught an average of 22.0 females and 0.3 males over 18 hours. In open areas, attraction of females to the volunteers showed a first peak shortly after sunrise between 07:30 and 08:00 h and a second, larger peak was observed around sunset between 19:00 and 21:30 h. At shady sites, the first peak was delayed to between 09:30 to 10:30 h and the second appeared earlier, between 18:00 to 19:30 h. The morning peak of female activity in shaded areas, averaging 15 landings per volunteer, was greater than the average of 9 landings per volunteer in open areas. In the evening, the average number of female landings per volunteer was 18 in open areas, while in shade it was nearly 27. Interestingly, in shaded areas there were also small numbers of male landings, 5 landings per volunteer, roughly at the same time as the peak of female activity. In open areas, landing of males was negligible (Fig 5A).

The attraction rate to a sugar source (traps baited with flowers of *P. juliflora*) in sunny and shaded areas also exhibited two peaks as did the search for a host (Fig 5B). In open areas, the first smaller peak of female landings was just after sunrise and the second followed sunset. In shaded areas, the first peak occurred between 11:30 h and 12:30 h while the second took place from around 16:30 h to 17:30 h. Sugar questing activity of males in the shady areas followed the pattern and duration of the females but the numbers caught were about a half. Only 2 males were caught in open sunny areas throughout the entire study period (Fig 5B).

### Comparing sugar feeding rates of *Ae. aegypti* in ‘sugar rich’ and ‘sugar poor’ habitats

Less than 20% of females and nearly 80% of males landing on human volunteers were sugar positive in both sugar rich and sugar poor environments. From resting sites within vegetation, 60-65% of ‘sugar rich’ site females or males were sugar positive and so were 39-40% of the females or males sampled from the ‘sugar poor’ site. Leaving mosquitoes alive in cages for six hours before testing decreased the proportion of sugar positive specimens by about a half (Fig 6A and 6B).

**Fig 6.**
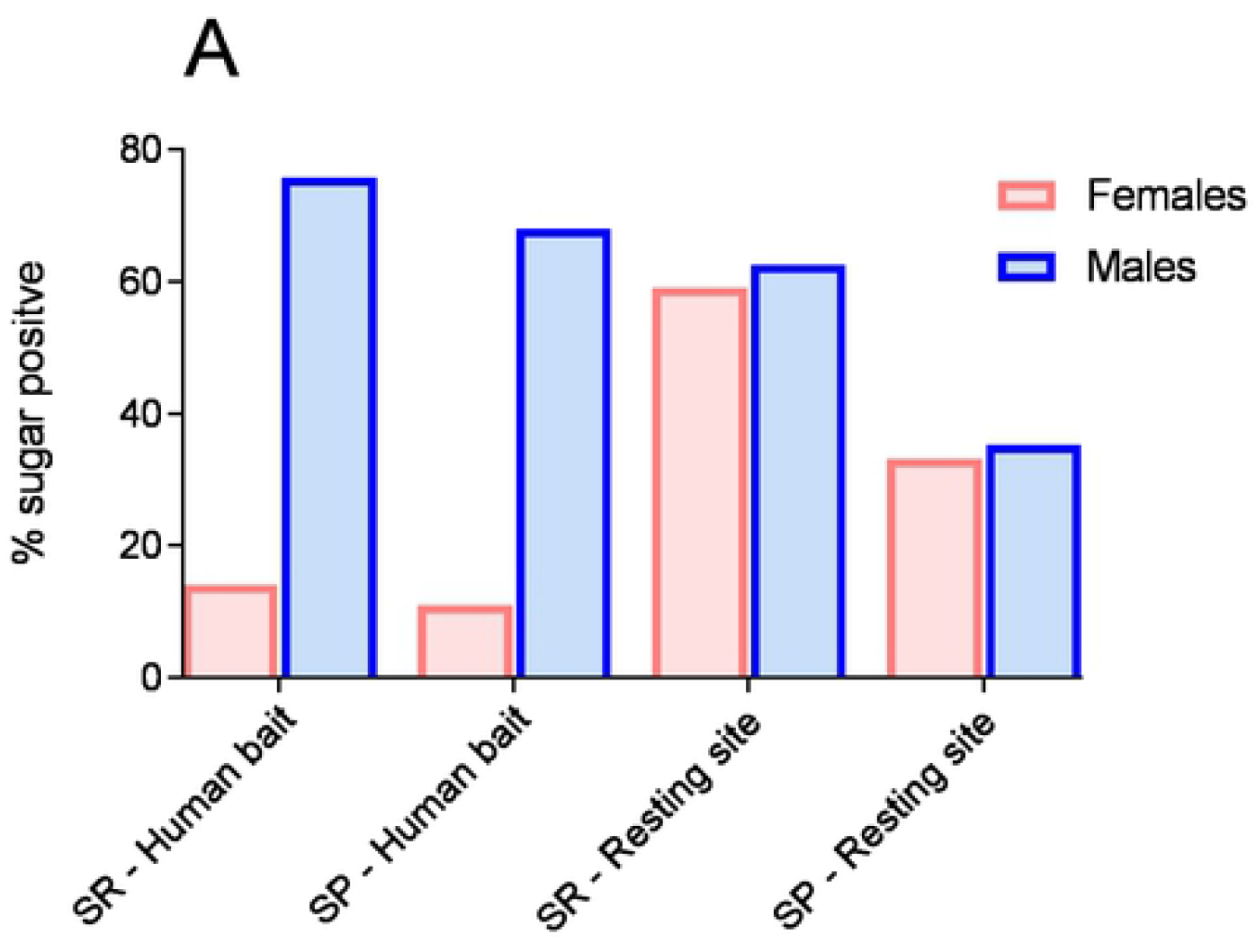
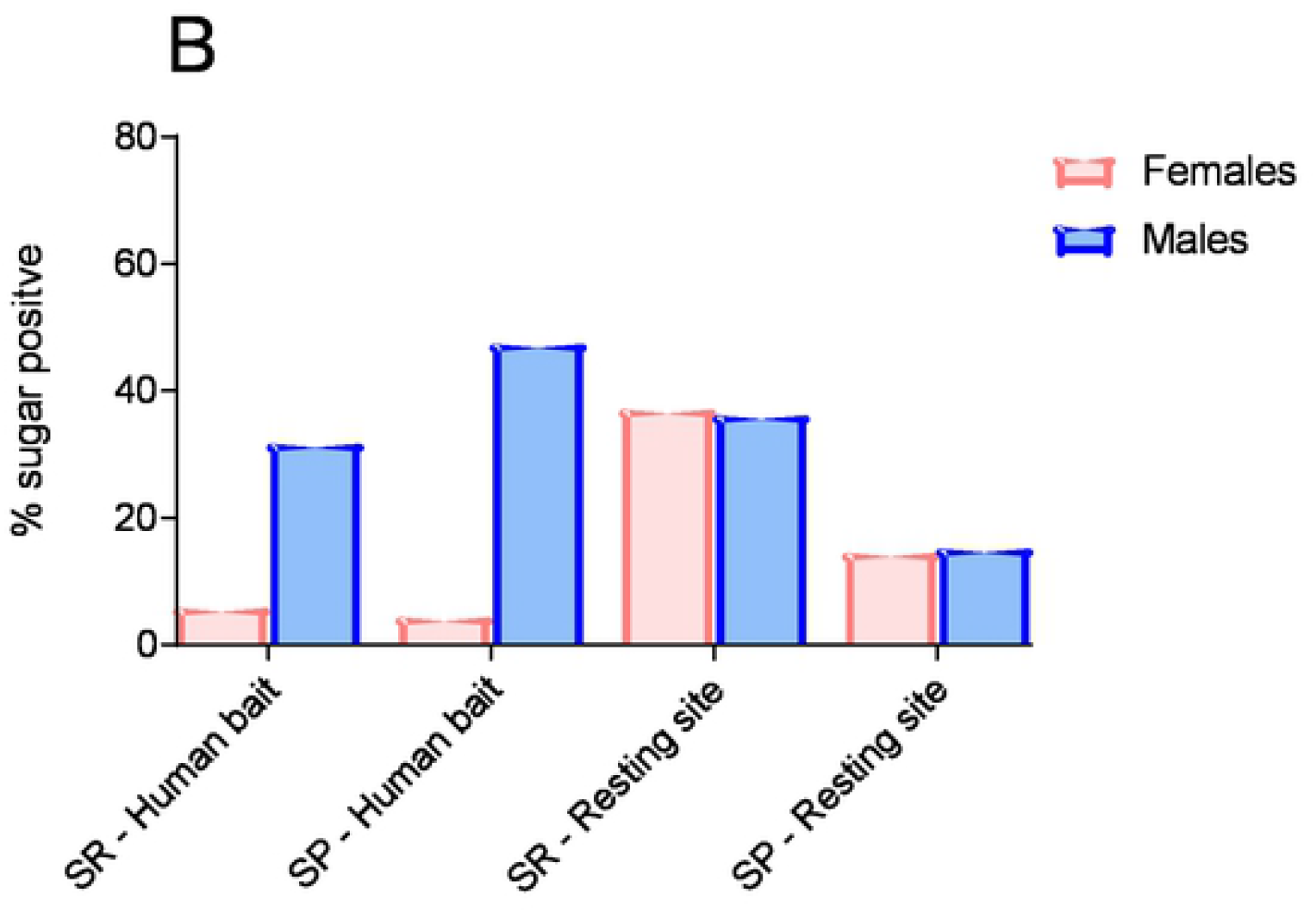
Percentage of sugar positive mosquitoes from the catch on volunteers and from resting habitat sweep-net catches at sugar rich (SR) and sugar poor (SP) sites. **(A)** Tested for sugar within 1 hour after collection and **(B)** 6 hours after collection.

Classification of *Ae. aegypti* females or males with different quantities of sugar in the gut showed that in ‘sugar rich’ environments, 60 to 90% of the sugar positive females caught on volunteers had mostly small (class I) sugar quantities in the gut (Table 3). Samples from ‘sugar rich’ resting sites, contained various sugar quantities in similar proportions (classes I to IV). A 6 hour delay in testing for sugar caused the sugar contents in the gut to degrade, and thus the reactions in the sugar testing to fall below classes III and IV. The sugar feeding status of mosquitoes caught at the ‘sugar poor’ sites, on volunteers or in resting habitats, followed the pattern observed for mosquitoes from the ‘sugar rich’ sites.

### ATSB experiments

The baseline for evaluating the impact of ATSB treatment was the density of female mosquitoes landing on volunteers at the control site, and the pre-treatment period at the experimental site. The mean daily catches on the 3 volunteers increased gradually at both ‘sugar rich’ and ‘sugar poor’ control sites. At the ‘sugar poor’ control site, the mean began at 5.46 ±0.43 females and was 32.93±2.89 on the last experimental day. At the ‘sugar rich’ control site, the initial mean increased from 12.27 ±1.20 to 65.89 ±5.55 females on the last day. ATSB treatment at both experimental sites on day 15 caused a drastic reduction in the numbers of *Ae. aegypti* females from about 5 days post-application until the end of the experiments (P < 0.001 days 16-50; Table 4). At the treated ‘sugar poor’ site, the mean of captured females on the last day was 0.29 ±0.10 and at the ‘sugar rich’ site it was 3.20 ±0.43 (Fig 7A and 7B, Table 4). ATSB significantly reduced mean numbers of landing / biting female *Ae. aegypti* at both ‘sugar rich’ and ‘sugar poor’ sites. At the sugar poor site, the mean catch pre-treatment on day 14 was 20.51 females and was reduced 70-fold to 0.29 on day 50. At the sugar rich site, the mean pre-treatment catch on day 14 was 32.46 and was reduced 10-fold to 3.20 on day 50 (Fig 7A and 7B, Table 4).

**Fig 7.**
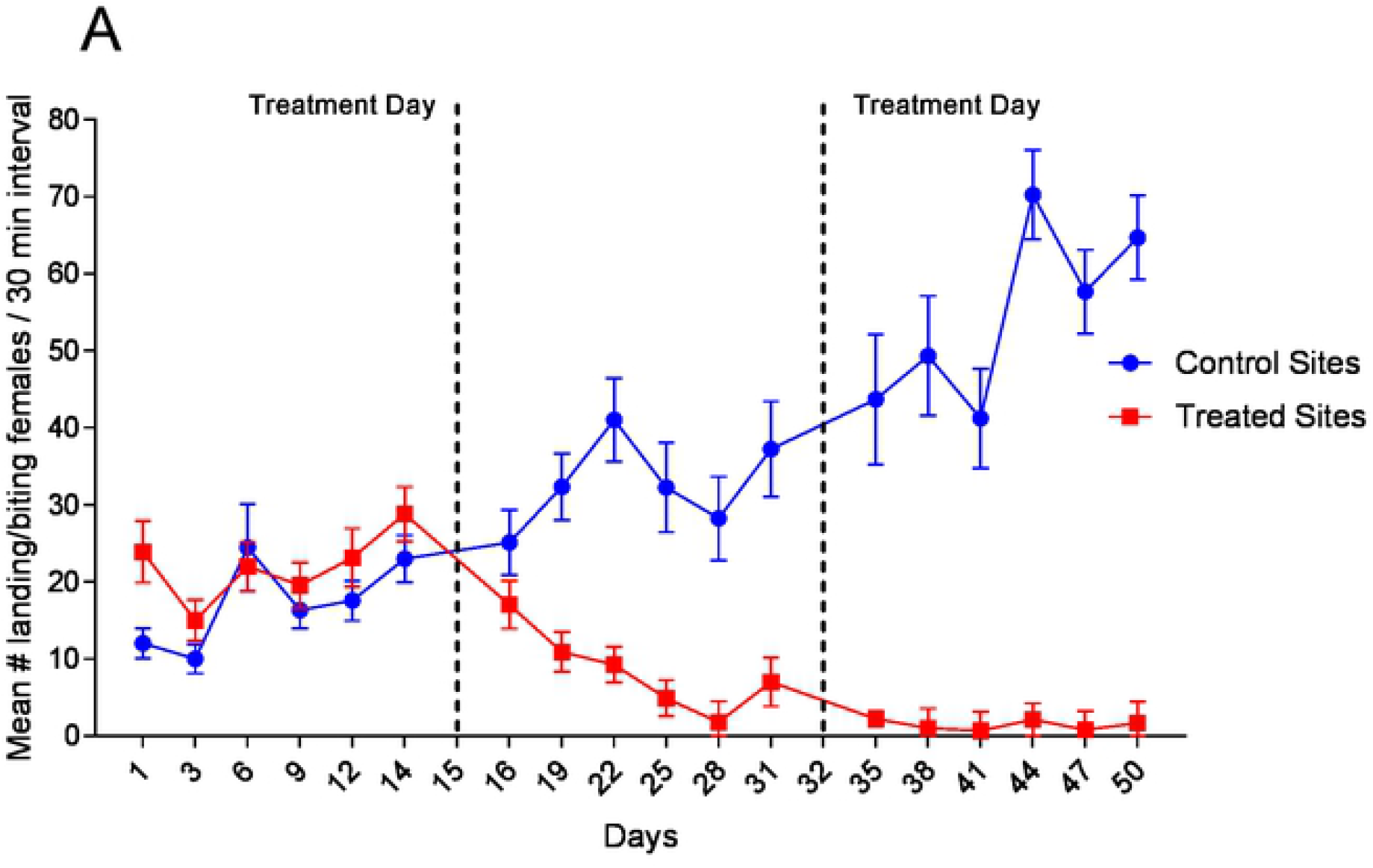
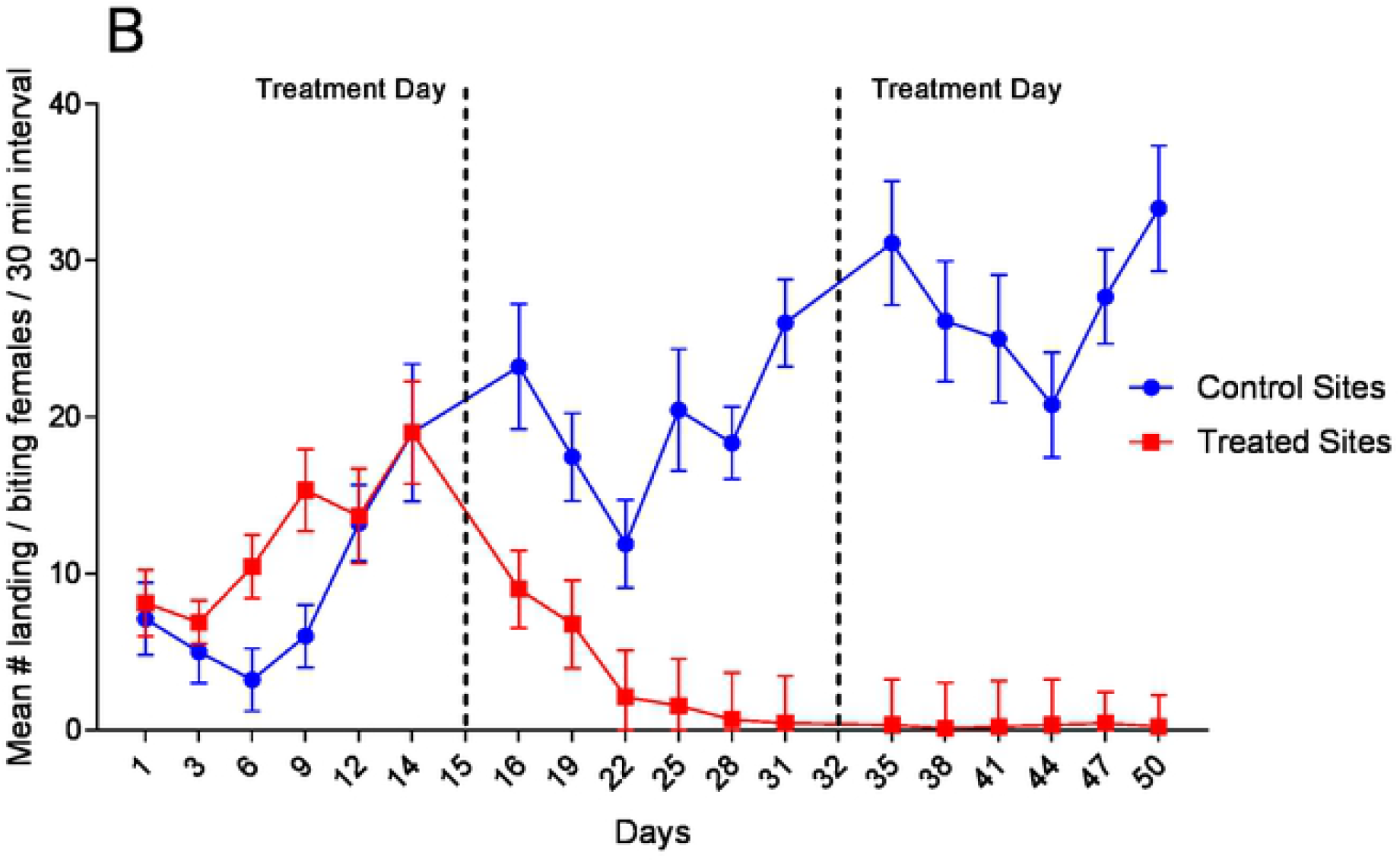
Reduction of *Ae. aegypti* female population following ATSB treatment. Results are shown as the reduction in mean number of landing/biting attempts per 30-minute interval for the treatment and control sites. (A) Sugar rich sites. (B) Sugar poor sites.

## Discussion

In this study, we investigated the feeding of *Ae. aegypti* in the laboratory and in the field on some natural sugar sources. We compared sugar feeding of the mosquitoes in urban ‘sugar poor’ and ‘sugar rich’ habitats, and then tested the potential of mosquito control by ATSB in the two different types of environments.

The 24 hour exposure to single, potential sugar sources demonstrated that the proportion of sugar positive *Ae. aegypti* was high in those offered flowers or fruit: 93.00% of females and 92.78% of males fed on *Acacia macrostachia*, 84.44% of females and 97.22% of males fed on *C. papaya*, 93.89% of females and 83.89% of males fed on *C. melo*, 81.67% of females and 84.44% of males fed on on *M. indica*, and 92.78% of females and 93.33% of males fed on *P. juliflora*. It is interesting to note that feeding rates for females (and for males) on some flowers was low: 15.56% of females fed on *B. glabra* and 1.67% of females fed on *N. glauca*. Seedpods (*P. reticulatum*) and sugarcane (*S. officinarum*) have tough outer coverings and therefore their sugar is apparently inaccessible to *Ae. aegypti* (Table 1). It is not surprising that mosquitoes died in less than a week on the sole diet of fairly inedible *B. glabra* while ~ 90% survived in a series maintained on the highly attractive nectar of *P. juliflora*. A high rate of feeding does not necessarily mean high quality of meals. In the survival experiment, there was no link between the 85.00% of females feeding on *G. gracilis* flowers within 24 hrs (Table 1) and the high (~ 50%) mortality in mosquitoes exposed to these flowers for 30 days (Fig 4A and 4B). In this context, it should be noted that the mosquitoes in the above experiments may have fed on a given source of sugar because there was no alternative. It should also be noted that in the tight space of 50 × 50 cm cages, contact between the flying mosquitoes and the offered diet is presumably inevitable, hence feeding is not necessarily the outcome of attraction.

The preference of sugar sources in nature is exhibited in the size of catches by baited traps which show the relative attraction compared to other tested baits and in competition with other environmental olfactory cues. An example of the difference between direct contact with a sugar source in cages and the performance of a sugar source as an attractant is *L. camara*. Overnight exposure to it in a cage resulted in 60.0 to 63.3% feeding (Table 1) but used as a bait in the field, the low catch amounted to a mean of 1.3% to 3.6% females per trap (Table 2A).

Such differences in attraction in laboratory versus field settings were noted and discussed in early studies [15,38,39]. Some of these attractive baits were also highly attractive to *An. gambiae* s.l., *An. sergentii, Ae. albopictus* and *Culex pipiens* [27,28,30,39] showing that mosquitoes of different genera and of both sexes are commonly attracted to specific emanations mostly to flowers of some plants. It would be interesting to see whether it is a developed adaptation that guides different mosquitoes to the richest sources of sugar in their environment.

The observation of peak times for searching for sugar meals, which are at dawn and dusk (Fig 5A and 5B), are useful for obtaining maximal information on the sugar-feeding status of *Ae. aegypti* populations. This is particularly important in view of the rapid digestion of sugar meals which may be misleading when mosquitoes are caught at the wrong time of the day or kept alive before being tested for sugar. For example, in catches on volunteers in sugar rich area only 60.32% blood fed mosquitoes had stage 1 (minimal quantities) sugar quantities in the gut. Examination of a similar series of mosquitoes caught in sugar rich areas, after 6 hrs delay, the proportion of stage 1 blood meals increased to 90.39%. In previous studies that estimated the digestion rates of mosquito species in the field, the half-life of fructose was approximately 7 to 10 hours [15,40,41].

To a certain degree, sugar meals inhibit the taking of blood and vice versa, both competing for space in the gut system [42, 43] and as a result, host seeking behaviour is terminated [44]. In Israel [34], it was shown that *Ae. albopictus* females that have fed on natural sugar sources or ATSB did not come to humans to blood feed. An opposite extreme case is *Ae. aegypti* that are adapted to domestic environments where sugar may be rare but there is an easy and unlimited access to blood [45]. This mosquito uses supplementary blood meals during the gonotrophic cycle as energy reserves [19] that convert blood into survival time in a ratio reported to be higher than that of nondomestic species [17]. In the domestic environment, females of this species seldom feed on sugar, but feral populations often do [45, 46]. In the laboratory, all sugar feeding ceases when blood-host stimuli are present, but without such stimuli, sugar feeding is frequent [47]. Thus, the option to take sugar is retained for the competitive advantage it affords under some circumstances. A similar strategy may be used by some anthropophilic *Anopheles* spp. [48]. Whether *Ae. aegypti* commonly, rarely, or never take sugar in nature remains controversial [48-51].

Theoretically, such variations can be interpreted as results of metabolic differences between mosquito subpopulations. Otherwise, assuming that producing energy from blood meals is general in *Ae. aegypti*, differences in the rates of blood and sugar feeding could be a response dictated by the environment that is, according to the relative abundance of sources.

It was also concluded that the enhanced blood-feeding capability among older sugar-deprived *An. gambiae* demonstrated the close association between sugar-feeding and blood-feeding behavior [52]. Our results similarly portray dependence of the blood feeding drive on sugar feeding. In catches on volunteers, the proportion of sugar positive females was similar in sugar poor (68.37% class I, Table 3) and in sugar rich areas (60.32% class I, Table 3). On the other hand, in sugar rich resting habitats, the quantity of class I sugar meals was 13.18% and 65.35% were of classes III and IV. Moreover, their proportion was about 40% greater than in the sugar poor resting habitats where 46.31% were of class I and 25.51% were of classes III and IV. These results indicate that following feeding on larger sugar quantities in the relatively lush vegetation of urban sugar rich habitats, mosquitoes were less interested in blood meals. Also, in the sugar poor and sugar rich areas, the effect of ATSB treatments were manifested at similarly rapid rates and the reduction of the mosquito population was to similar levels. In other words, mosquitoes responded equally when the sugar bait (ATSB) was uniformly offered in both habitats. In both experiments, the catch on volunteers and the results of ATSB treatment, it appears that the intensity of the search for a host blood meal depends on the prevalence or scarcity of sugar meals in different types of *Ae. aegypti* urban habitats.

Attractive Toxic Sugar Bait (ATSB) treatment applied in the sugar poor and sugar rich areas caused a drastic reduction in *Ae. aegypti* approaches to volunteers in both environments. The initial effect of ATSB was apparently somewhat delayed since there was a massive supply of newly emerged mosquitoes. Later, high female mortality reduced the oviposition and the *Ae. aegypti* populations collapsed almost completely. At the sugar rich sites, the daily approaches of *Ae. aegypti* to the volunteers decreased from an average of 28.0 before treatment to 1.7 landings / bites a week later (Fig 7A). At the sugar poor sites, an average of 19.0 landings was reduced to 0.44 in the first week and these levels remained similarly low until the end of the experiment (Fig 7B). The drastic effect of ATSB is comparable to the results obtained in Israel, Florida, Morocco and Mali [28, 30, 33, 53-55].

Since sugar is the only food source for male mosquitoes [15], ATSB should be highly effective against male *Ae. aegypti* but should also affect survival and fecundity of females as well. Our data demonstrates that ATSB treatment can be highly effective against *Ae. aegypti* in both sugar rich and sugar poor environments. ATSB was also highly effective against *Ae. albopictus* in Mali [34] and it may be a significant treatment against both species, particularly if their distribution overlaps. Generally, it appears from this study that ATSB is a new promising tool for the control of *Ae. aegypti*.

## Acknowledgements

We dedicate this work to Fatoumata Sissoko who passed away suddenly after the completion of this work. We also wish to thank the participating communities of Bamako in Mali. This work, titled “ICEMR-Mali: Multidisciplinary research for malaria control and prevention in West Africa”, was supported by NIH NIAID grant number U19 AI129387 awarded to JB. The funders had no role in study design, data collection and analysis, decision to publish, or preparation of the manuscript. The authors declare that no competing interests exist in relation to this project.

